# Beyond microRNAs: Analysis of chimeric reads characterises the diverse targetome of AGO2-mediated regulation

**DOI:** 10.1101/2023.08.24.554582

**Authors:** Vaclav Hejret, Nandan Mysore Varadarajan, Eva Klimentova, Katarina Gresova, Ilektra-Chara Giassa, Stepanka Vanacova, Panagiotis Alexiou

## Abstract

Argonaute proteins are instrumental in regulating RNA stability and translation. AGO2, the major mammalian Argonaute protein, is known to primarily associate with microRNAs, a family of small RNA ‘driver’ sequences, and identifies its targets primarily via a ‘seed’ mediated partial complementarity process Despite numerous studies, a definitive experimental dataset of AGO2 ‘driver’-’target’ interactions remains elusive. Our study employs two experimental methods - AGO2 CLASH and AGO2 eCLIP, to generate thousands of AGO2 target sites verified by chimeric reads. These chimeric reads contain both the AGO2 loaded small RNA ‘driver’ and the target sequence, providing a robust resource for modeling AGO2 binding preferences. Our novel analysis pipeline reveals thousands of AGO2 target sites driven by microRNAs and a significant number of AGO2 ‘drivers’ derived from fragments of other small RNAs such as tRNAs, YRNAs, snoRNAs, rRNAs, and more. We utilize convolutional neural networks to train machine learning models that accurately predict the binding potential for each ‘driver’ class and experimentally validate several interactions. In conclusion, our comprehensive analysis of the AGO2 targetome broadens our understanding of its ‘driver’ repertoire and potential function in development and disease. Moreover, we offer practical bioinformatic tools for future experiments and the prediction of AGO2 targets. All data and code from this study are freely available at https://github.com/ML-Bioinfo-CEITEC/HybriDetector/

## Introduction

The AGO clade of the Argonaute protein family is a widely conserved set of proteins with regulatory functions. In mammals, four AGO proteins (AGO1-4) are known to primarily associate with small non-coding RNA molecules called microRNAs (miRNAs) to form ribonucleoprotein complexes that include an AGO protein loaded with a miRNA ‘driver’ sequence (Fig.1A). This ‘driver’ sequence leads this ribonucleic complex to target specific RNAs, regulating their levels via mechanisms of translational silencing and/or degradation [1]. This miRNA ‘driver’ mediated targeting is a central regulation mechanism in metazoans, flies, and other animals, earning miRNAs the apt term ‘Sculptors of the Transcriptome’ [2]. Out of the four mammalian AGO proteins, solely Ago2 single loss causes embryonic lethality in murine models [3, 4] and so can be considered as the most important member of a partially redundant family of proteins.

**Fig. 1.**
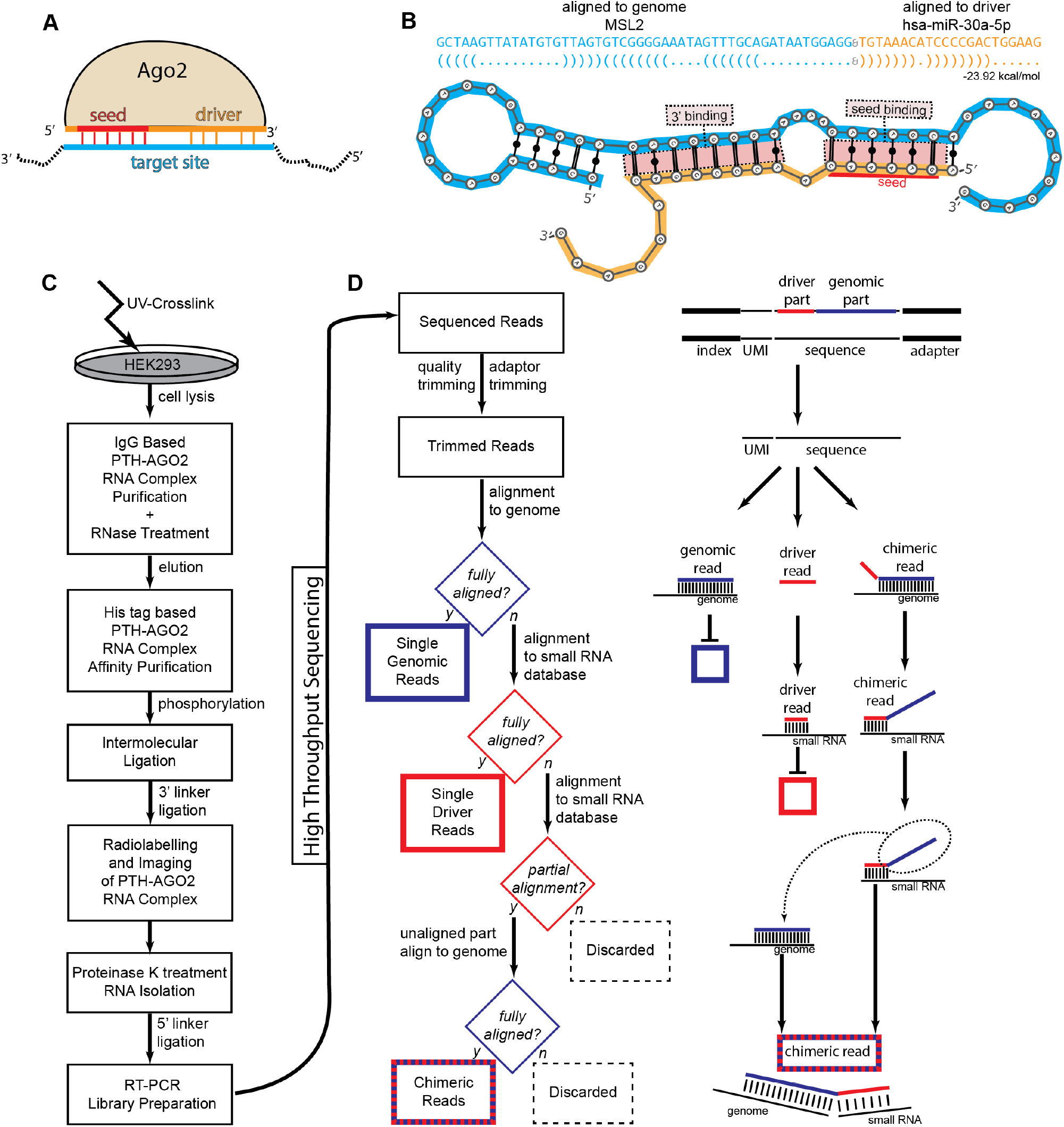
(A) Schematic of Ago2 loaded with a short RNA driver sequence, binding to a target site using a seed driven approach. (B) Schematic of a chimeric read containing fragments of the small RNA driver sequence and the target site. (C) Experimental outline of the CLASH technique. (D) Outline of the bioinformatic pipeline for identification of single genomic, single small RNA, and chimeric reads.

After AGO2 is loaded with a miRNA ‘driver’ (AGO2:miRNA) it can use partial complementarity between the ‘driver’ to identify ‘target’ sequences on other RNA molecules. Unlike plant miRNAs that have fully complementary targets [5], in mammals most known AGO2:miRNA binding sites show partial complementarity focused on a ‘seed’ region located at the 5’end of the ‘driver’ sequence. A ‘canonical seed’ sequence denotes a fully Watson-Crick complementary stretch of at least 6 nucleotides starting at the second position from the 5’ end of the miRNA ‘driver’. Further binding outside the seed area could have a stabilising effect on the interaction [6]. However, functional interactions not mediated by a ‘canonical seed’ have been known since the early days of miRNA targeting research in worms [7] and mammals [8]. The exact rules of AGO2:miRNA target recognition remain unknown, but several approximations have been produced to date as crucial parts of miRNA target prediction programs.

Most such methods use a two-step approach. In the first step, putative binding sites are identified using either a seed-based or a ‘cofolding’-based approach. The seed-based approach tries to identify canonical or almost canonical seed sequences, and weigh them based on categories of binding [9]. The cofolding approach uses methods that calculate the minimal energy of folding between the miRNA and the putative target sequence (Fig.1B), using this as a prioritization and weighing technique [10]. As a second step, various features of the identified putative binding sites are combined into an overall prediction of the probability of a specific AGO2:miRNA to repress a target RNA as a whole. When the first generation of miRNA target prediction programs was evaluated on independent benchmarks of mRNA translational inhibition, methods using seed heuristics consistently outperformed ‘co-folding’ based methods [11]. The seed-based rules of early miRNA target prediction programs were based on a handful of known and experimentally validated interactions.

In 2011 a new experimental method, termed CLASH (crosslinking, ligation, and sequencing of hybrids) was developed [12] which uses a ligation step between the ‘driver’ and ‘target’ sequences and can produce ‘chimeric’ reads containing both sides of the AGO:driver:target interaction. Such an experiment was performed for the AGO1 protein, which surprisingly revealed that only 40% of the identified miRNA binding sites contained a canonical seed in human cells [13]. This finding reopened the question of binding rule identification beyond the seed, even though non-canonical seed contribution to functional targeting may be hard to estimate in bulk [9].

Given the abundance of bona fide non-seed interactions, target prediction methods using only ‘canonical’ seed as the prime determinant of binding, may be missing out on predictive sensitivity, by ignoring a large number of interactions. To compound the issue, the systematic pro-seed bias feeds back into the ‘experimental validation’ loop, amplifying its impact on future development.

Another important finding from the AGO1-CLASH experiment is that miRNAs are not the only ‘drivers’ loaded on the AGO1 protein. Sequencing of small RNA fragments associated with AGO proteins identified various other types of RNAs, such as fragments of tRNAs, snoRNAs, vaultRNAs and others [14]. Fragments of tRNAs (tRFs) were later shown to associate with AGO proteins [15, 16] and confer post-transcriptional silencing regulation to their targets, in a manner similar to miRNAs [17–19]. Recently, human ribosomal RNA fragments were identified by computational meta-analysis of AGO1 immunoprecipitation and CLASH experiments as potentially functional drivers of AGO1 targeting [19].

In this paper we present the first dataset of AGO2-CLASH experimental data, a bioinformatics pipeline for driver:target identification that takes into account non-miRNA ‘drivers’, as well as trained and tested machine learning models based on Convolutional Neural Networks that can accurately identify AGO2 binding sites, outperforming both ‘seed’ and ‘co-fold’ based methods. Recently, a novel experimental method for miRNA chimeric read sequencing was developed, called miR eCLIP [20]. We have also produced a complementary dataset based on this method (AGO2-eCLIP), which we use to validate our findings from the AGO2-CLASH method.

## Methods

### Human cell culture

The Human Embryonic Kidney 293 T-Rex FlpIn (HEK293T) with inducible expression of hAGO2-PTH was given to us by the Tollervey lab [21]. Cells were cultured in DMEM supplemented with 10% FBS in an atmosphere of 5% CO2, 37°C.

### AGO2-CLASH

We followed the protocol established by (Helwak et al. 2013) with some modifications. All chimeras produced by AGO2-CLASH can be found in Supp. Table ST1. See details in Supplementary Methods.

### AGO2-eCLIP

Two experiments following the miR-eCLIP methodology [20] were performed by Eclipse Bioinnovations using Eclipse AGO2 IP antibodies against AGO2 on HEK293xT cells. No modifications were made to the standard miR-eCLIP methodology for this experiment. All chimeras produced by AGO2-eCLIP can be found in Supp. Table ST2.

Chimeras of two driver sequences TR1: ‘TCCGGCTCGAAGGACCA’, and TR2: ‘TCCCGGGTTTCGGCACC’ were optionally enriched in the AGO2-eCLIP experiment. For purposes of reporting chimeric read abundances, any driver sequence with edit distance score < 5 to any of the two sequences was removed from consideration.

### Luciferase assays

Validations of predicted interactions were carried out using the dual luciferase reporter system psiCHECK2 plasmid and Promega Dual Luciferase Reporter Assay Kit. An exact methodology of the luciferase assay can be found in Supplementary Methods.

### AntimiR assays and Quant-seq

High-throughput analysis of effects of inhibiting targeted miRNA was carried out using AntimiRs for 24hrs against hsa-miR-320a, and hsa-miR-484 at final concentration of 30nM. The global transcriptome effects of each miRNA’s inhibition were measured using Quant-Seq. Detailed methodology can be found in Supplementary Methods.

### Chimeric Read Annotation Pipeline (HybriDetector)

Chimeric reads produced by the CLASH or miR eCLIP protocols consist of two distinct interacting RNA molecules, partially digested and connected by intermolecular ligation at one end. However, chimeric reads are only a small fraction of the sequenced library, as single ‘driver’ and single ‘target’ reads can be found. The goal of HybriDetector is to separate these types of reads, and annotate the ‘drivers’ and ‘targets’ as accurately as possible. A detailed technical overview of the pipeline can be found in Supplementary Methods. The HybriDetector pipeline itself is freely available at https://github.com/ML-Bioinfo-CEITEC/HybriDetector/

### Convolutional Neural Network

We trained Convolutional Neural Networks (CNNs) consisting of six layered blocks, each composed of a Convolutional layer, leaky ReLU, batch normalization, pooling, and a dropout layer. The output of the last dropout layer is flattened and connected to a dense neural network. The last layer is formed of a single neuron with a sigmoid activation function that outputs the probability of driver:target site binding. A detailed technical overview of the model and training scheme can be found in Supplementary Methods. Full trained models, and all code used for training and evaluation is freely available at https://github.com/ML-Bioinfo-CEITEC/HybriDetector/tree/main/ML

## Results

### Identification of chimeric ‘driver’-’target’ interactions

We have performed the AGO2 CLASH experiment in three replicates by using the HEK293T–hAGO2-PTH cell line, following the CLASH protocol [12, 13] with minor modifications (Fig.1C). We sequenced the CLASH samples using NextSeq 500 from Illumina, yielding a total of 730,731,281 reads with a maximum read length of 75bp. We have designed and implemented a bioinformatic pipeline, HybriDetector, for the identification of chimeric reads from CLASH and similar experiments, with a focus on the annotation of reads involving several types of non-miRNA ‘driver’ sequences. We separate three categories of reads [‘single-driver’, ‘single-genomic’, ‘chimeric’] through a series of alignments on the genome or small RNA sequence databases (Fig. 1D) miRNA, tRNA, rRNA, snoRNA, YRNA or vaultRNA. Reads strongly mapping on any of these non-coding RNA annotation databases, are defined as ‘single-driver’ sequences. Finally, reads that have one part mapping on the masked genome, and one on the non-coding RNA annotations are considered as ‘chimeric’. As expected, ‘driver’ reads tend to be shorter than 25nt, while ‘genomic’ and ‘chimeric’ reads are closer to the full length of 75nt (Fig 2A). From these chimeric reads, duplicated records are removed using Unique Molecular Identifiers and post-process filters are applied, such as no mismatch allowed in alignments, support of the chimeric alignment target by overlapping single reads alignment, alignment lengths of individual chimeric read parts, and others (full list reported in Suppl. Methods).

**Fig. 2.**
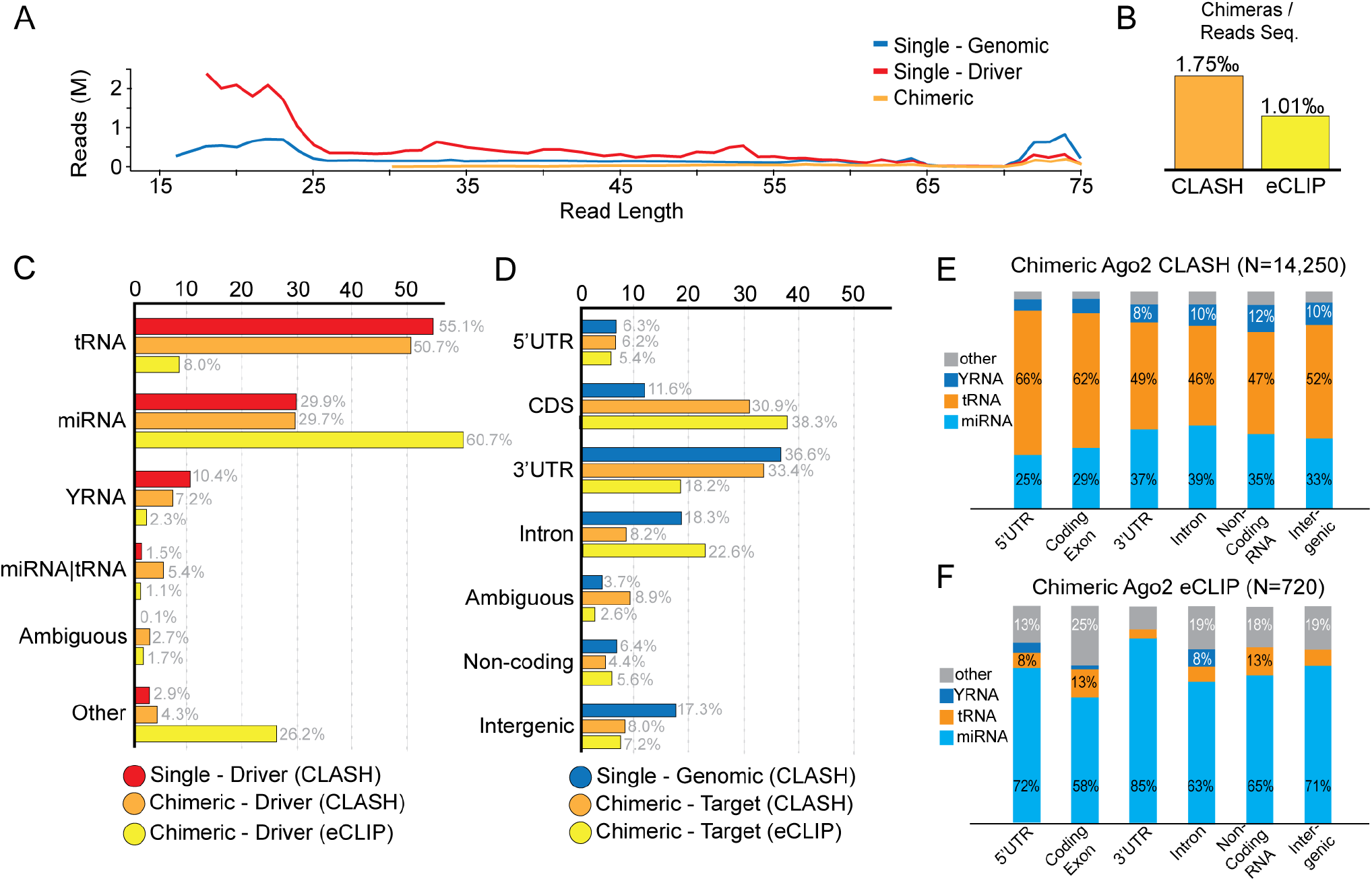
(A) Read length distribution for sequencing reads annotated as genomic, driver, and chimeric. (B) Fraction of high confidence chimeric interactions per thousand reads for AGO2-CLASH and AGO2-eCLIP experiments. (C) Distribution of identified driver sequences on driver databases. (D) Distribution of genomic target sequences on genic element annotations. (E) Distribution of AGO2-CLASH chimeric reads on driver databases and genic annotation. (F) Distribution of AGO2-eCLIP chimeric reads on driver databases and genic annotations.

Chimeric reads coming from the same ‘driver’ and mapping at the same ‘target’, allowing for minor sequence discrepancies, are collapsed into a single representative ‘chimeric interaction’. We thus obtain a very strict list of 14,205 high confidence ‘chimeric interactions’ (Suppl. Table TS1) which are the dataset used for all further analyses.

When mapping AGO2-CLASH reads against the miRNA reference, we discovered that the miRNA with the most annotated chimeric reads was miR-4286. The sequence of miR-4286 (ACCCCACUCCUGGUACC) is also found in tRNA-Leu-TAA, also known as tRF-3009a (ACCCCACUCCUGGUACC**A**). This tRF has been associated with disease such as lupus (Geng et al., 2021) and cardiomyocyte response to glucose (Zhao et al., 2023). In turn, miR-4286 has been associated with cancer (Komina et al., 2016; Ho et al., 2022) and as a biomarker for acute coronary disease (Shen et al., 2021). None of these studies considers the sequence similarity between miR-4286 and tRF-3009. We examined CLIP data of the miRNA biogenesis factor Drosha on HEK293T cells [22] and discovered that in this cell line, miR-4286 does not appear to have any Drosha activity, while the tRNA-Leu-TAA loci seem to be covered in a manner similar to other miRNAs, such as let-7a (Suppl. Fig. S3). Out of caution, we have marked these interactions, and others with similar characteristics as ‘ambiguous’ and removed them from any downstream analysis.

### Comparison of AGO2-CLASH and miR eCLIP results

We produced miR eCLIP (AGO2-eCLIP) libraries from HEK293T cells without exogenous AGO2 induction, and analyzed them through the HybriDetector pipeline. We did sequence the AGO2-CLASH libraries much deeper than the AGO2-eCLIP libraries, resulting in 14,250 chimeric interactions (1.75‰ sequenced reads) for AGO2-CLASH versus 720 chimeric interactions (1.01‰ sequenced reads) for miR eCLIP (Fig. 2B)

The AGO2-CLASH method gave consistently more tRNA derived single drivers (56.1%), as well as chimeric drivers (50.7%) against miRNA derived single drivers (29.9%), and chimeric drivers (29.7%). This is consistent with analysis of the AGO1-CLASH dataset (43.9% tRNA chimeric drivers, 31.4% miRNA chimeric drivers). However, the AGO2-eCLIP library gave more chimeric miRNA derived drivers (60.7%) against tRNA ones (8.0%), when enriched sequences were removed (Fig. 2C). Since the AGO2-eCLIP experiment does not include an AGO2 induction step, we can report that tRNA derived drivers do get loaded on AGO2 under physiological conditions, however, the overexpression of AGO in the CLASH experiment tends to overestimate their abundance.

Both the AGO2-CLASH and the AGO2-eCLIP methods showed a distribution of target sites mostly between coding sequence exons, and 3’UTRs (Fig. 2D). Interestingly, the AGO2-eCLIP data showed approximately one quarter of chimeric targets on introns (22.6%). Intronic targets for miRNAs have also been identified using a method similar to CLASH based on a pan-AGO antibody against all 4 Argonaute proteins in mouse and human cell lines (36% intronic chimeras) [23]. It is interesting to note, that intronic targets are effectively ignored by all miRNA target prediction methods. Cross-referencing ‘driver’ provenance and genic annotation of the target site, we notice no major difference between the distributions of different ‘driver’ types within the AGO2-CLASH distributions (Fig 2E). However, for AGO2-eCLIP (Fig. 2F) we notice that 85% of chimeric interactions mapping on 3’UTRs appear to come from miRNA drivers, with that number dropping to 58% when considering coding exons.

### Experimental validation of single targets

We decided to experimentally validate if individual chimeric interactions identified in our AGO2-CLASH experiment could indeed modulate translation. We transfected HEK293T cells with driver mimics and a dual luciferase reporter construct containing their corresponding target sequences within the 3’ UTR of Renilla. We chose a variety of driver-target chimeras from high-confidence chimeric reads of diverse origins.

Out of 17 unique driver-target pairs tested, twelve demonstrated that the chimeric small RNA could suppress Renilla expression that contained the respective target mRNA sequence in the 3’UTR (Fig. 3E). Intriguingly, alongside the noted miRNAs (let-7, miR-320a, and miR-484), we discerned the functionality of small RNAs originating from noncoding RNAs (two tRNAs and Y1 RNA) on several mRNA chimeric targets.

**Fig. 3.**
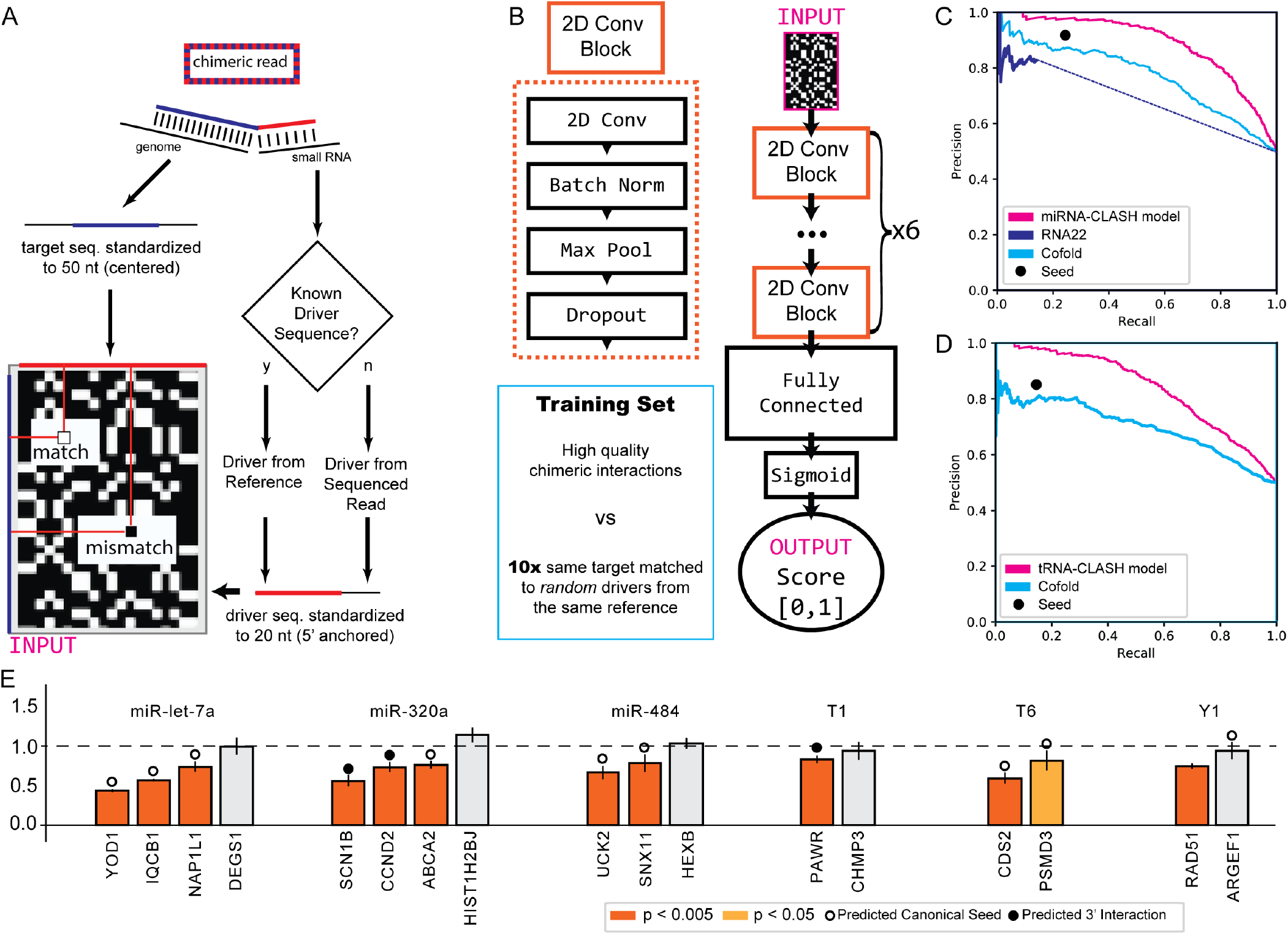
(A) Process of chimeric read representation as 2D alignment matrix. (B) Architecture of the Convolutional Neural Network used for training binding prediction models, consisting of three 2D Convolutional blocks followed by a fully connected network. All models were trained on 1:10 imbalanced datasets derived from high quality chimeric interactions. (C) Precision-Recall curve of miRNA trained CNN model against the state of the art, evaluated on left-out balanced high quality chimeric interactions. (D) Precision-Recall curve of tRNA trained CNN model against the state of the art, evaluated on left-out balanced high quality chimeric interactions. (E) Luciferase assay validation of selected chimeric interactions. Interactions with a predicted canonical seed (circle), predicted 3’ interaction but no seed (dot), and no clear interaction were select-ed. R/F ratios below 1.0 denote efficient downregulation upon transfection, orange and yellow bars showing significant downregulation within replicates using Student t-test.

Most pairs (9) featured a canonical seed sequence match between the miRNA and its target (from the 2nd to the 8th nucleotide), and three pairs exhibited a match of five or more nucleotides at the 3’ end sequence terminus. Among the five pairs that showed no inhibitory effect, only one pair had a canonical miR-seed match (miRY1-ARGEF1), whereas the remaining four lacked either a canonical or a 3’ terminal extensive match.

In summary, these experiments demonstrated our ability to reveal significant miRNA-mRNA target pairs and the functional potential of newly discovered small RNAs, which are derived from noncoding RNAs, along with their prospective mRNA targets.

### Chimeras correlate with mRNA level changes after miRNA knockdown

In order to further validate the identified chimeric binding sites, we employed anti-miRs to inhibit two highly expressed miRNAs: miR-320a and miR-484. We subsequently employed Quant-Seq to monitor global alterations in mRNA expression levels.

In the case of miR-320a, we determined that mRNAs encompassing at least one high-confidence chimeric interaction exhibited significant upregulation after 24 hours of anti-miR transfection (Wilcoxon Rank Sum Test p=2e-06), compared to expressed mRNAs without chimeric interactions. We observed a similar trend in the miR-484 inhibition experiment (Wilcoxon Rank Sum Test p=0.0022), although the effect was less pronounced (Suppl. Fig. S4)

It’s crucial to acknowledge that chimeric interactions are infrequent occurrences, resulting in a limited sample size for our study. As a consequence, it’s entirely feasible that many of the ‘control’ mRNAs are actual targets of the inhibited miRNAs. Therefore, the observed trend should be interpreted as a cautious estimate of the effect.

These two datasets of gene expression changes after anti-miR mediated inhibition will also make a great asset for future research into the effects of miRNA repression at the gene level.

### Binding site prediction using a CNN model

Given that the majority of recognized AGO2:miRNA interactions lack a standard seed sequence and existing co-folding methods have shown limited capacity to prioritize functional binding [11], there is a pressing need for novel computational approaches to identify and prioritize targets. We previously established miRBind, a Convolutional Neural Networks (CNNs)-based technique trained on AGO1-CLASH data, which exhibited superior performance in predicting AGO1 binding sites compared to existing methodologies [24].

Building on insights garnered from the aforementioned study, we designed a CNN capable of learning binding interactions from a two-dimensional representation of the Watson-Crick binding potential of the ‘driver’ and ‘target’ sequences (Fig. 3A). This CNN is composed of six 2D Convolutional Blocks, each housing a 2D convolution, Batch Normalization, Max Pooling, and Dropout layers. Subsequently, the output is routed through two fully connected layers, and a sigmoid activation function, culminating in a final prediction between 0 and 1, indicating the likelihood of the two sequences binding under AGO2 conditions (Fig. 3B). Additional information regarding the training and optimization of the CNN is available in the supplementary methods.

We benchmarked our models against the canonical six nucleotide ‘seed’ measure, the RNA co-fold energy serving as a binding score, and RNA22, the only available miRNA target prediction software providing binding scores solely from sequence (Loher and Rigoutsos, 2012). In each instance, our models surpassed the standard models in the classification task for both miRNA (Fig. 3C) and tRNA (Fig. 3D) chimeras. The performance of the miRNA trained model was assessed using left-out samples of its corresponding type from AGO2-CLASH, as well as against miRNA derived chimeric interactions from the AGO2-eCLIP experiment where its performance matches that of the canonical seed measure (Supplementary Fig. S5).

Compared to the AGO2-CLASH dataset, the AGO2-eCLIP dataset features a higher enrichment of canonical seed chimeric interactions. When evaluated against the independent AGO2-eCLIP dataset, our miRNA model exhibited comparable precision/recall to the canonical seed, while demonstrating greater versatility in identifying less constrained targets (Supplementary Fig. S5). These results validate the utility of our trained model for a wholly independent experiment, one potentially enriched in legitimate endogenous targets, given that it does not involve an AGO2 induction step.

## Discussion

Since the discovery of AGO proteins, canonical targeting by miRNA drivers on 3’UTR targets has been the main paradigm of AGO function [25], despite some early known non-canonical seed interactions [26]. This bias has been driven by bioinformatic target prediction programs that prioritize canonical seed interactions. To break this cycle, we executed the first ever Ago2-CLASH experiment and AGO2-eCLIP technique to identify a wide array of Ago2 interactions, confirming the role of non-seed targets, and non-miRNA AGO2 ‘drivers’. However, our analysis also indicated that CLASH techniques overestimate the number of non-miRNA ‘drivers’. Further study is needed to understand the propensity of AGO to load more tRNA and other non-miRNA short RNAs when overexpressed.

Keeping into consideration that miRNAs are not the only potential ‘drivers’ for Ago2, we have developed a bioinformatic HybriDetector, a publicly available pipeline that can be used to extract and disambiguate binding sites based on several types of potential ‘driver’ types, including miRNAs, snoRNAs, tRNA fragments, and other non-coding RNAs. Our findings also indicate that sequence similarity between different ‘driver’ types can mislead researchers and that machine learning models can improve binding site prediction over canonical seed heuristic.

We were able to validate the functionality of several driver:target pairs identified in our AGO2-CLASH analysis that included both annotated miRNAs as well as newly identified small RNAs derived from noncoding RNAs (tRNAs and Y-RNA). The target pairs that did not show an effect in the reporter system were mostly those where we did not detect any potential for miRNA seed match base pairing. It is possible that such chimeras could result from a background ligations caused by endogenous RNA ligase as previously reported in *C. elegans* [27]. Recent studies identified an RNA ligase in human cells which could confer this ligation [28].

Our results show that AGO2-eCLIP is easier to perform experimentally than AGO2-CLASH and may become the dominant method. An additional advantage of AGO2-eCLIP is the use of endogenous AGO2 which seems to make a significant difference on the type of ‘drivers’ loaded by AGO2.

Our expectation of an increase in the number of such experiments is based on the AGO2-eCLIP method’s simplicity and effectiveness, combined with the significant impact of using endogenous AGO2 on the type of ‘drivers’ loaded. This increase will correspondingly generate a larger and more diverse dataset of ‘driver’ and ‘target’ interactions for researchers to study and understand.

This larger dataset will not only improve the predictive accuracy of our machine learning models but also enable them to capture a wider array of interactions and nuances. This improved understanding will lead to the refinement of our current bioinformatic pipeline, helping us to better disambiguate binding sites and potentially identify novel ‘driver’ types.

Furthermore, an increased quantity and variety of experimental data will allow the exploration of AGO2’s role in a broader context. It could open avenues to understanding differential AGO2 behavior across various cell types, developmental stages, and disease conditions.

Additionally, an expanded dataset could also aid in unveiling the rules that guide the loading of specific ‘drivers’ onto AGO2. It might also help investigate the factors influencing the specificity and effectiveness of AGO2’s interaction with different ‘drivers’ and ‘targets’, thereby revealing new aspects of post-transcriptional regulation and AGO2’s role in cellular functions.

We anticipate that the potential widespread adoption of the AGO2-eCLIP method will bring about a surge in experimental data, further advancing our understanding of small RNA biology, AGO2 function, and RNA-target interactions. This will ultimately enrich the resources available to the small RNA targeting community and foster the development of increasingly refined target prediction algorithms.

Recent advancements in machine learning have enabled us to develop highly precise sequence models that can predict the potential binding between a ‘driver’ and ‘target’ sequence, using AGO2-CLASH data. These models serve as a compelling alternative to the traditional seed-based heuristics commonly used in miRNA target prediction programs for initial filtering. Utilizing a convolutional neural network approach, we’ve successfully outperformed the conventional seed heuristic and the commonly used minimal energy of co-folding score. Though the limited quantity of available chimeric interactions restricts the size of the model that can be trained, the continuous influx of experimental data promises to enhance these models’ accuracy by allowing them to learn more comprehensively the binding rules for different ‘driver’ classes. Our models are publicly accessible and can serve as a first-step alternative to the seed in miRNA or tRNA mediated AGO2 targeting for researchers using target prediction programs.

An important future area of study will involve deciphering the deep learning models to extract human-readable rules. Understanding how these models achieve their superior accuracy compared to the simpler seed heuristic will provide a significant contribution to the small RNA binding community. In this study, we’ve treated the trained models as ‘black boxes’, examining the types of interactions they’ve learned to accurately identify. The miRNA and tRF models appear to have learned determinants beyond seed binding, as they assign higher scores to positive interactions without a canonical seed, over negative interactions with a canonical seed. Interestingly, when cross-evaluated, these models lose accuracy, indicating that they’ve learned at least partially different determinants beyond the seed. Notably, our models are designed not to see the actual ‘driver’ or ‘target’ nucleotide sequences; instead, we represent the interaction as a 2D matrix of potential Watson-Crick binding. This approach prevents our models from learning the exact sequences of ‘drivers’ or ‘targets’, ensuring their versatility and applicability to any new ‘driver’ sequence, irrespective of its presence in our training set.

To conclude, we have produced novel chimeric datasets for small RNA binding mediated by AGO2 in HEK293 cells using two complementary experimental techniques. We have developed a bioinformatic pipeline for the detection of chimeric reads that takes into account the difficulties of using multiple small RNA references. Finally, we have trained state of the art machine learning models to detect the potential for binding between AGO2 loaded with miRNAs or tRFs and its targets.

We anticipate that this study will pave the way for a new generation of target prediction algorithms. These algorithms will capitalize on the steadily increasing volume of chimeric reads, using them for training and improving the accuracy of binding interaction predictions. This approach represents a significant leap forward in the field, promising enhanced precision and effectiveness in RNA-targeting applications. Ultimately, we hope our findings contribute to a deeper understanding of small RNA binding mechanisms, informing future research and applications in the field.

## Acknowledgements

We would like to thank A. Helwak for the AGO2-PTH cell line, AH and Ales Obrdlik for helpful suggestions about the CLASH protocol. We thank Leona Svajdova and Karolina Vavrouskova for excellent technical support. We acknowledge the CF Genomics supported by the NCMG research infrastructure (LM2018132 funded by MEYS CR) and CF Bioinformatics CEITEC MU (LM2023067 funded by MEYS CR) for their support with obtaining scientific data presented in this paper.

## Author Contributions

PA and SV planned the project. PA and ICG had oversight of the bioinformatic aspects, SV and NMV carried out wet lab experiments, VH developed the chimeric analysis pipeline, EK and KG developed the CNN method. All authors wrote and edited the manuscript.

