## Supplementary Materials and Methods for "Beyond microRNAs: Analysis of chimeric reads characterises the diverse targetome of AGO2-mediated regulation"

^1^Central European Institute of Technology, Masaryk University, Brno, 62500, Czech Republic ^2^National Centre for Biomolecular Research, Faculty of Science, Masaryk University, 62500 Brno, Czech Republic ^3^Department of Applied Biomedical Science, Faculty of Health Sciences, University of Malta, MSD 2080 Msida, Malta ^4^Centre for Molecular Medicine & Biobanking, University of Malta, MSD 2080 Msida, Malta

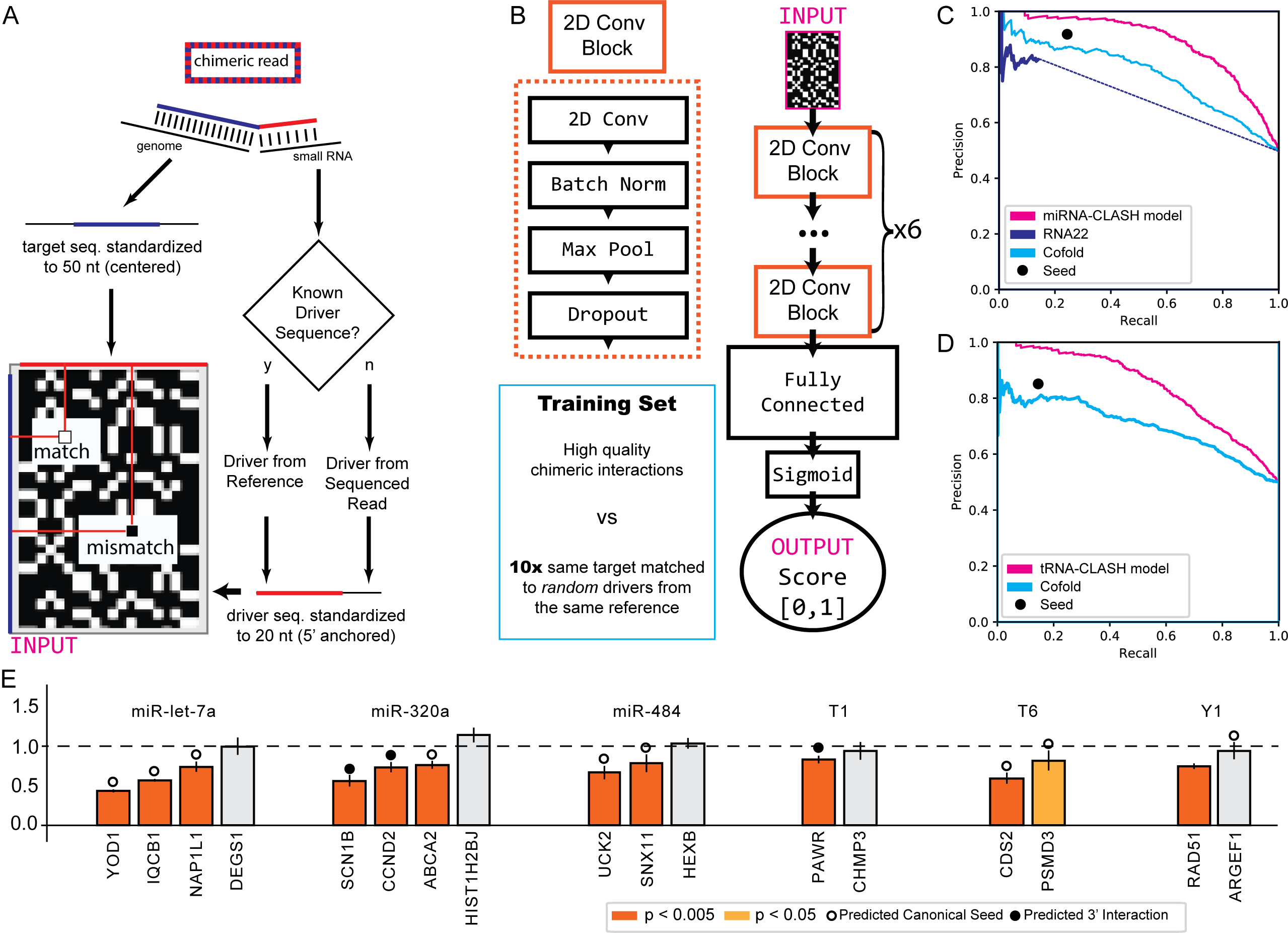

### Discussion

Since the discovery of AGO proteins, canonical targeting by miRNA drivers on 3’UTR targets has been the main paradigm of AGO function [25], despite some early known non-canonical seed interactions [26]. This bias has been driven by bioinformatic target prediction programs that prioritize canonical seed interactions. To break this cycle, we executed the first ever Ago2-CLASH experiment and AGO2-eCLIP technique to identify a wide array of Ago2 interactions, confirming the role of non-seed targets, and non-miRNA AGO2 'drivers'. However, our analysis also indicated that CLASH techniques overestimate the number of non-miRNA ‘drivers’. Further study is needed to understand the propensity of AGO to load more tRNA and other non-miRNA short RNAs when overexpressed.

### Declarations

**Ethics approval and consent to participate**
Not applicable
**Consent for publication**
Not applicable **Availability of data and materials**
All data and code from this study are freely available at <https://github.com/ML-Bioinfo-CEITEC/HybriDetector/> **Competing interests**
The authors declare that they have no competing interests. **Funding**
This work has been supported by the Czech Science foundation grants (No 19-10976Y to PA and 20-19617S and 23-07372S to SV), the institutional support CEITEC 2020 (LQ1601), and the HORIZON-WIDERA-2022 grant BioGeMT (ID: 101086768) to PA.
**Authors' contributions**
PA and SV planned the project. PA and ICG had oversight of the bioinformatic aspects, SV and NMV carried out wet lab experiments, VH developed the chimeric analysis pipeline, EK and KG developed the CNN method. All authors wrote and edited the manuscript.
**Acknowledgements**
We would like to thank A. Helwak for the AGO2-PTH cell line, AH and Ales Obrdlik for helpful suggestions about the CLASH protocol. We thank Leona Svajdova and Karolina Vavrouskova for excellent technical support. We acknowledge the CF Genomics supported by the NCMG research infrastructure (LM2018132 funded by MEYS CR) and CF Bioinformatics CEITEC MU (LM2023067 funded by MEYS CR) for their support with obtaining scientific data presented in this paper.

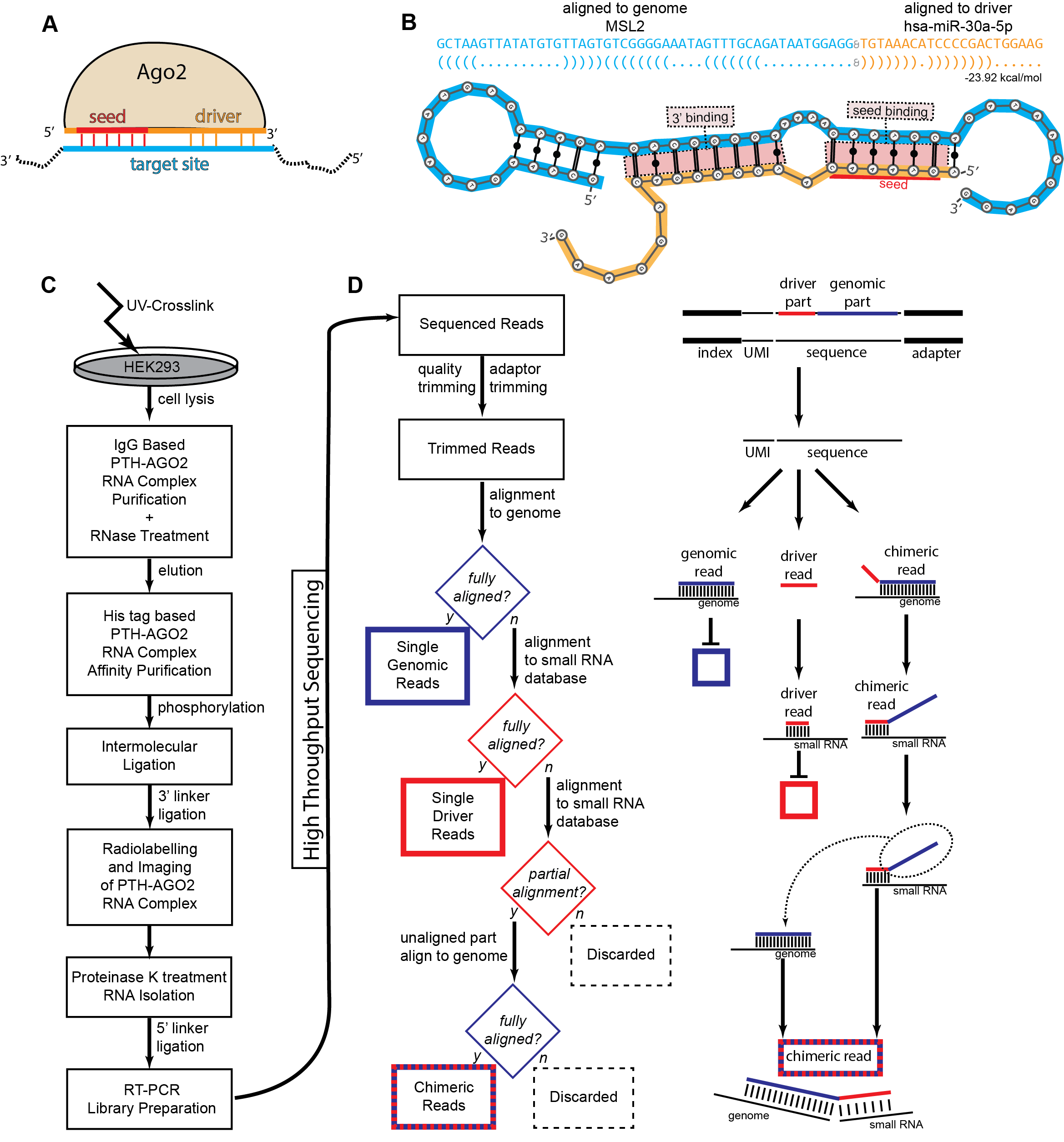
Fig. 1 (A) Schematic of Ago2 loaded with a short RNA driver sequence, binding to a target site using a seed driven approach. (B) Schematic of a chimeric read containing fragments of the small RNA driver sequence and the target site. (C) Experimental outline of the CLASH technique. (D) Outline of the bioinformatic pipeline for identification of single genomic, single small RNA, and chimeric reads.

Fig. 2 (A) Read length distribution for sequencing reads annotated as genomic, driver, and chimeric. (B) Fraction of high confidence chimeric interactions per thousand reads for AGO2-CLASH and AGO2-eCLIP experiments. (C) Distribution of identified driver sequences on driver databases. (D) Distribution of genomic target sequences on genic element annotations. (E) Distribution of AGO2-CLASH chimeric reads on driver databases and genic annotation. (F) Distribution of AGO2-eCLIP chimeric reads on driver databases and genic annotations.
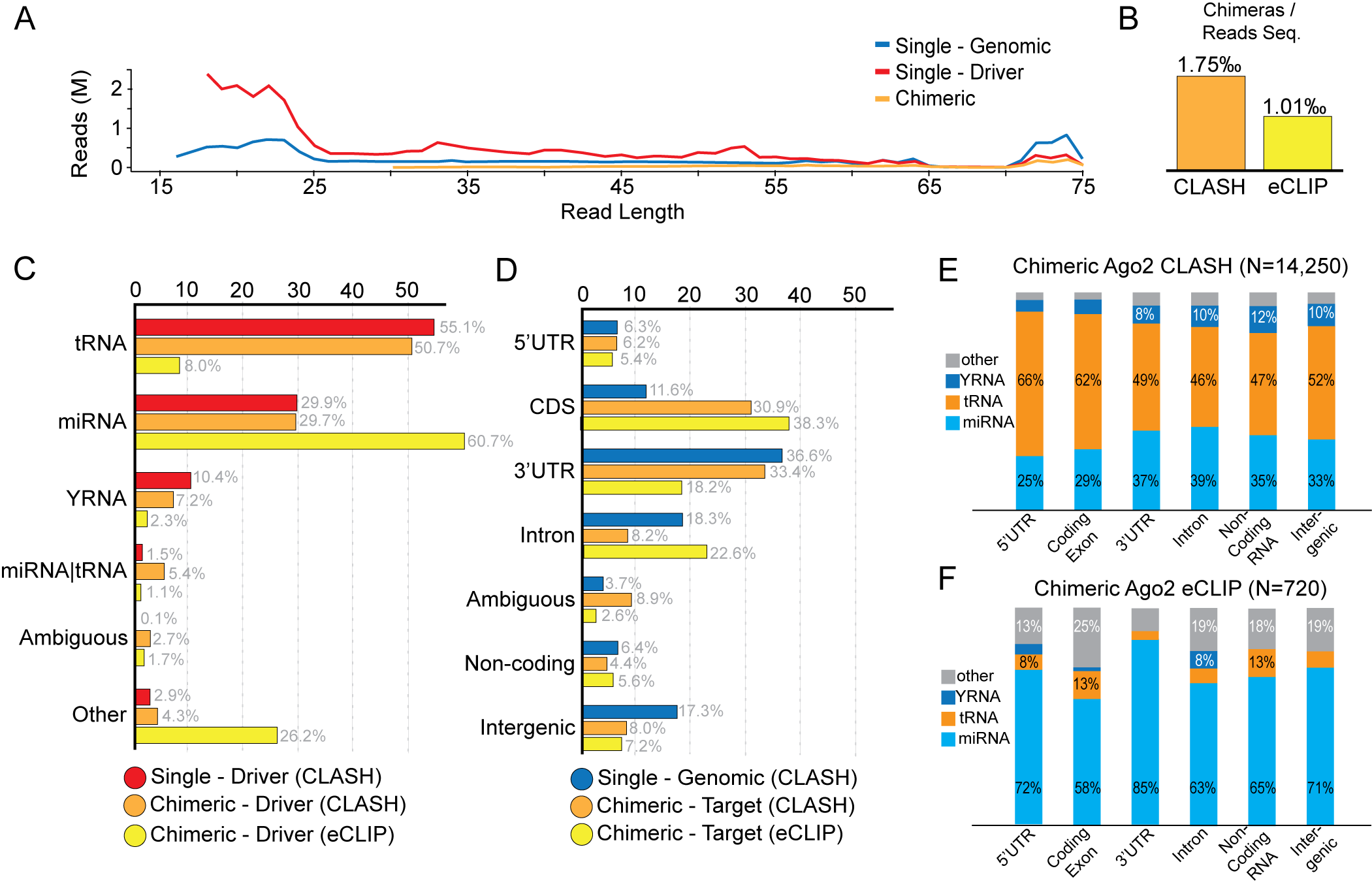

Fig. 3 (A) Process of chimeric read representation as 2D alignment matrix. (B) Architecture of the Convolutional Neural Network used for training binding prediction models, consisting of three 2D Convolutional blocks followed by a fully connected network. All models were trained on 1:10 imbalanced datasets derived from high quality chimeric interactions. (C) Precision-Recall curve of miRNA trained CNN model against the state of the art, evaluated on left-out balanced high quality chimeric interactions. (D) Precision-Recall curve of tRNA trained CNN model against the state of the art, evaluated on left-out balanced high quality chimeric interactions. (E) Luciferase assay validation of selected chimeric interactions. Interactions with a predicted canonical seed (circle), predicted 3’ interaction but no seed (dot), and no clear interaction were select-ed. R/F ratios below 1.0 denote efficient downregulation upon transfection, orange and yellow bars showing significant downregulation within replicates using Student t-test.
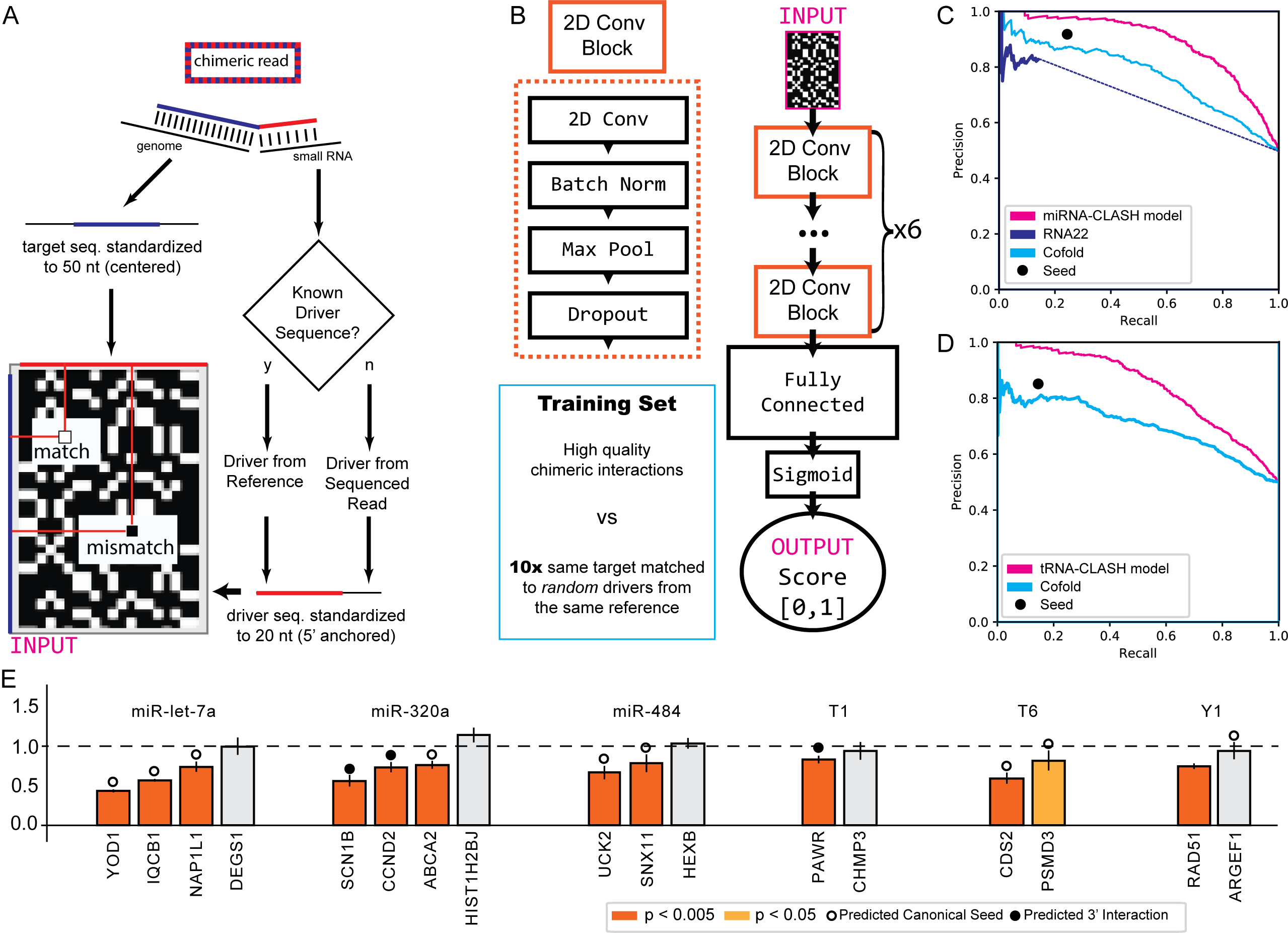
